## Supplementary Fig. 2 for "Epithelial fusion is mediated by a partial epithelial-mesenchymal transition"

A

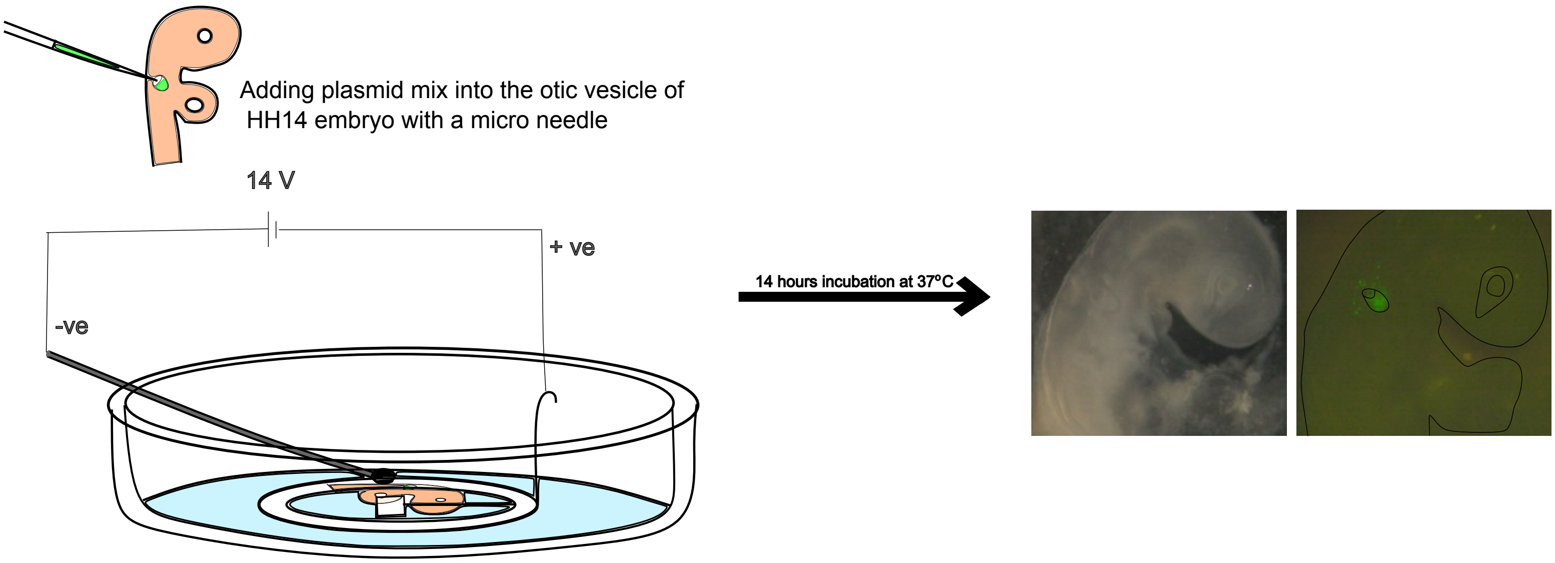


**Schematic for electroporation**

The embryo is placed in a dish connected to electrodes. The electrolyte used is ringer's solution and the voltage source is a square wave generator.
