## Supplementary figures and images for "Epithelial fusion is mediated by a partial epithelial-mesenchymal transition"

### Supplementary Fig. 1A

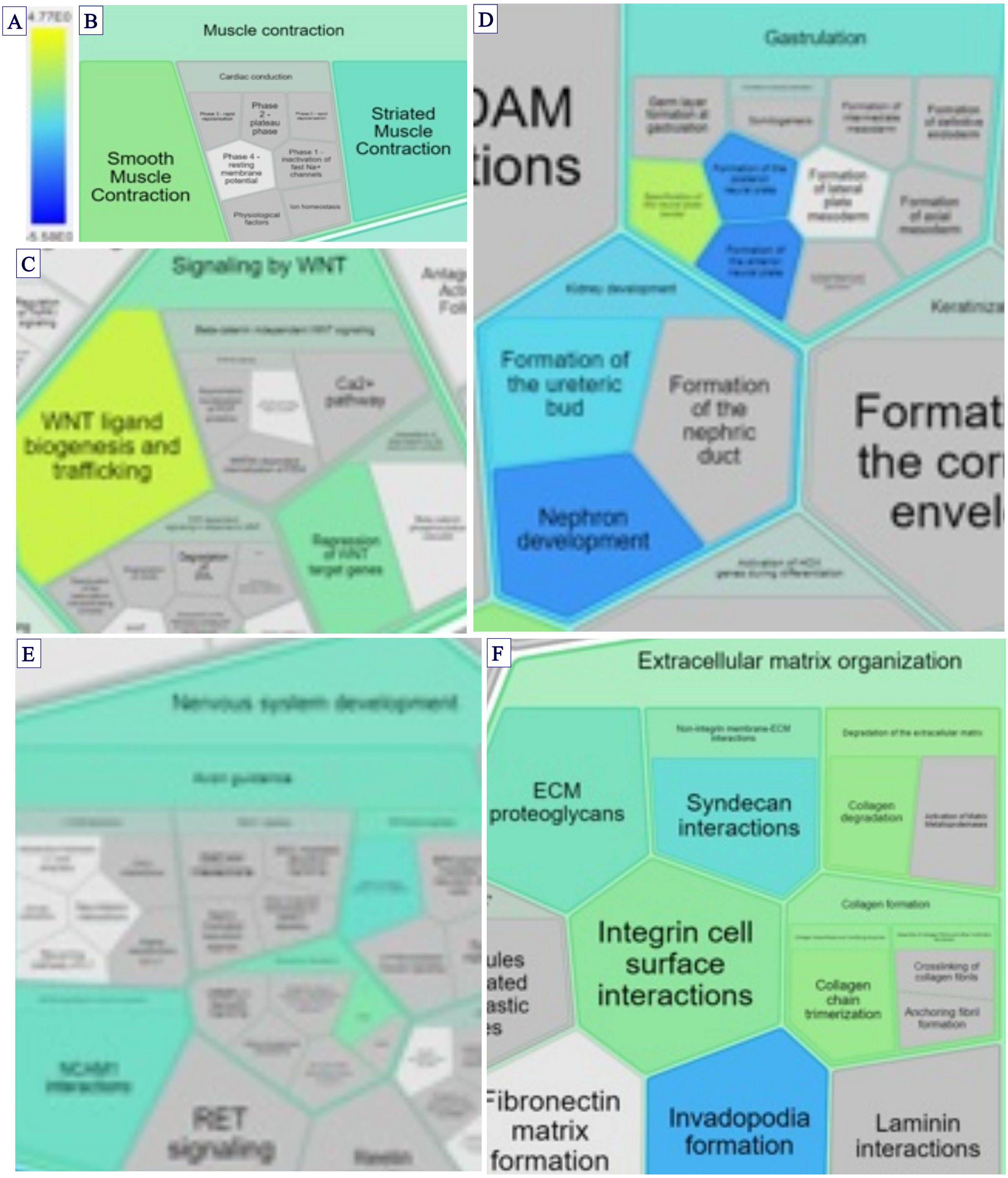

### Supplementary Fig. 1B

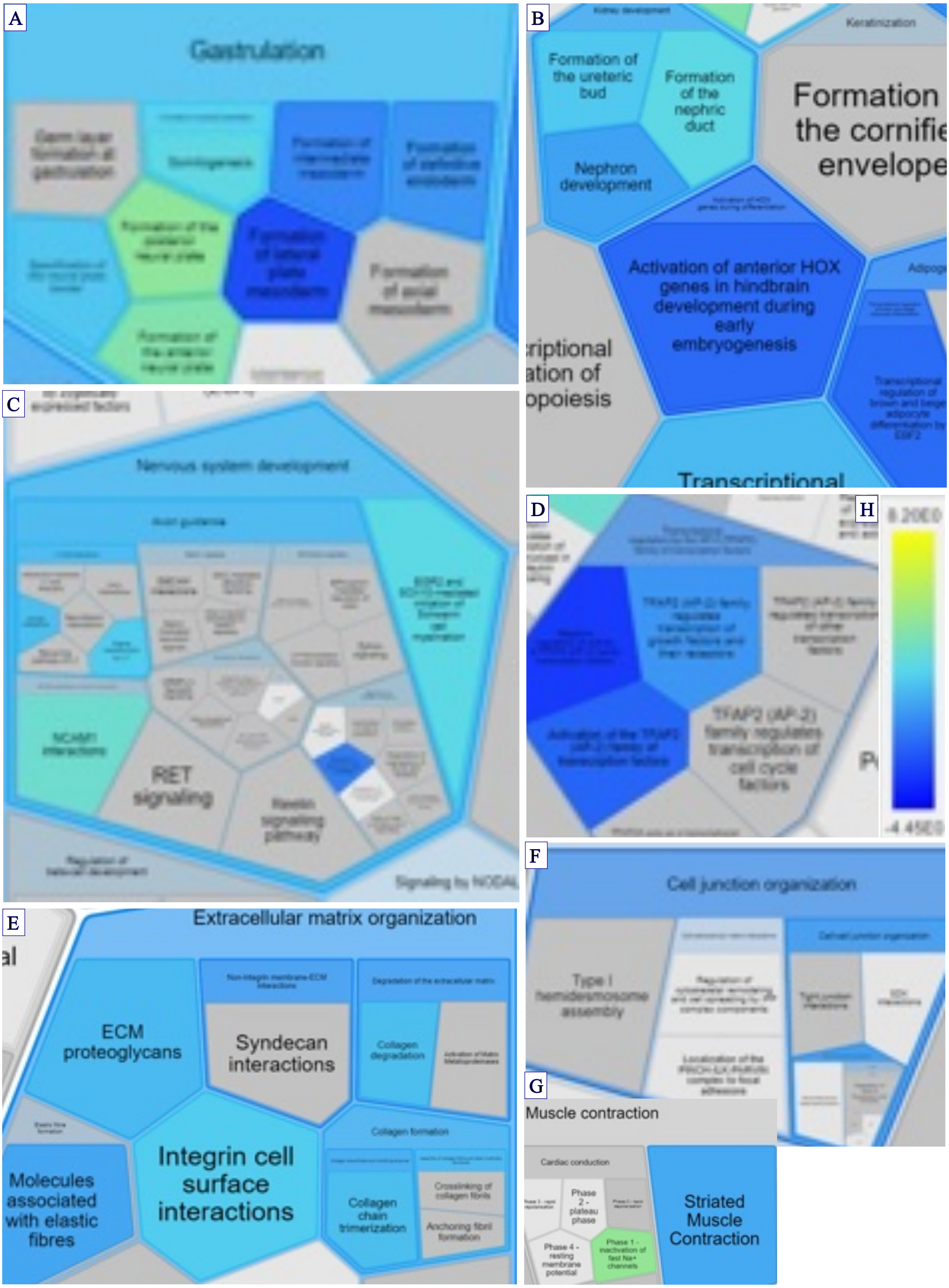
